# A quantitative autonomous bioluminescence reporter system with a wide dynamic range for Plant Synthetic Biology

**DOI:** 10.1101/2023.02.20.529214

**Authors:** Camilo Calvache, Marta Vazquez-Vilar, Elena Moreno-Giménez, Diego Orzaez

## Abstract

Engineered autonomous bioluminescence (EAB) offers many potential applications in Plant Synthetic Biology, notably as *in vivo* reporter system. Current EAB reporter configurations are limited for quantitative applications due to low dynamic range. We reconfigured the *Neonothopanus nambi* fungal bioluminescence (NeoLuc) pathway to serve as a high-throughput and inexpensive reporter for quantitative analysis of gene expression. We showed that by configuring the first committed step in the pathway (HispS) as the transcriptional entry point instead of the fungal luciferase, the dynamic range of the output increased dramatically, equaling that of the FLuc/RLuc reporter, and outperforming it in high throughput capacity. Furthermore, the inclusion of an enhanced GFP as normalizer allowed transient ratiometric measurements in *N. benthamiana*. Fast and rich datasets generated by the NeoLuc/eGFP system enabled us to undertake the optimization of new challenging synthetic gene circuits, including a complex agrochemical/optogenetic dual input switch for tight control of engineered metabolic pathways.

## INTRODUCTION

Plant Synthetic Biology (PSB) aims to create plants with augmented capacities by engineering new metabolic pathways and robust synthetic gene circuits (SGCs) operating new-to-nature genetic programs. The development of SGCs can be speeded up by adopting the recursive Design-Build-Test-Learn (DBTL) framework^1^. Quantitative reporter systems are essential in the testing steps of the DBTL cycle^2^. Reporter genes act as final circuit actuators, generating easily scoreable readouts of fluorescence, luminescence or absorbance that proxy the transcriptional activity generated by the upstream elements in the circuit, facilitating the acquisition of numeric values that feed mathematical models^3,4^. In plants, engineering through DBTL cycles is restricted by the lack of experimental platforms that support SGC testing in a fast and high throughput manner. The dual-ratiometric Firefly luciferase (FLuc) / Renilla luciferase (RLuc) reporter (FLuc/RLuc) is, ought to its wide dynamic range, ease of use, and low endogenous activity, the current golden standard for SGC evaluation in plants^5,6^. In this system, the promoter transactivation of FLuc serves as a transcriptional input, which is proportionally converted into luminescence readouts by FLuc enzyme, maintaining a high dynamic range. In parallel, the luminescence of a constitutively-expressed RLuc serves as an output normalizer. Despite its advantages, FLuc/RLuc involves laborious protein extraction, and most importantly, it requires the external addition of expensive luciferase substrates, limiting its use as a throughput data-rich reporter system in PSB.

The recently discovered bioluminescence pathway from the fungus *Neonothopanus nambi* has opened new possibilities for development of optimal reporter systems in PSB. The fungal bioluminescence pathway (FBP) is a cyclic metabolic route comprising four enzymes with no functional homologs in plants (see Fig. 1a). The fungus employs caffeic acid as the substrate of the first committed step in the pathway. Caffeic acid is a common intermediary metabolite in plants; therefore, plant cells can use their own metabolic pool of caffeic acid to feed the FBP autonomously, as it was recently demonstrated^7^. The third enzyme in the pathway, LUZ, is the actual luciferase, converting the metabolite 3-hydroxyhispidin into caffeylpyruvic acid and releasing a photon in the reaction. The facility for detection and the fact that there is no need to supply exogenously luciferin make FBP a promising reporter for plants^8^. However, to serve as a robust quantitative system in PSB, several improvements and adaptations are required. One of the main caveats lays in the low dynamic range that results from linking circuit transcriptional outputs to the expression of the fungal luciferase gene. Furthermore, in the absence of a second reporter gene, the variations in transfection efficiency cannot be accounted for.

**Figure 1.**
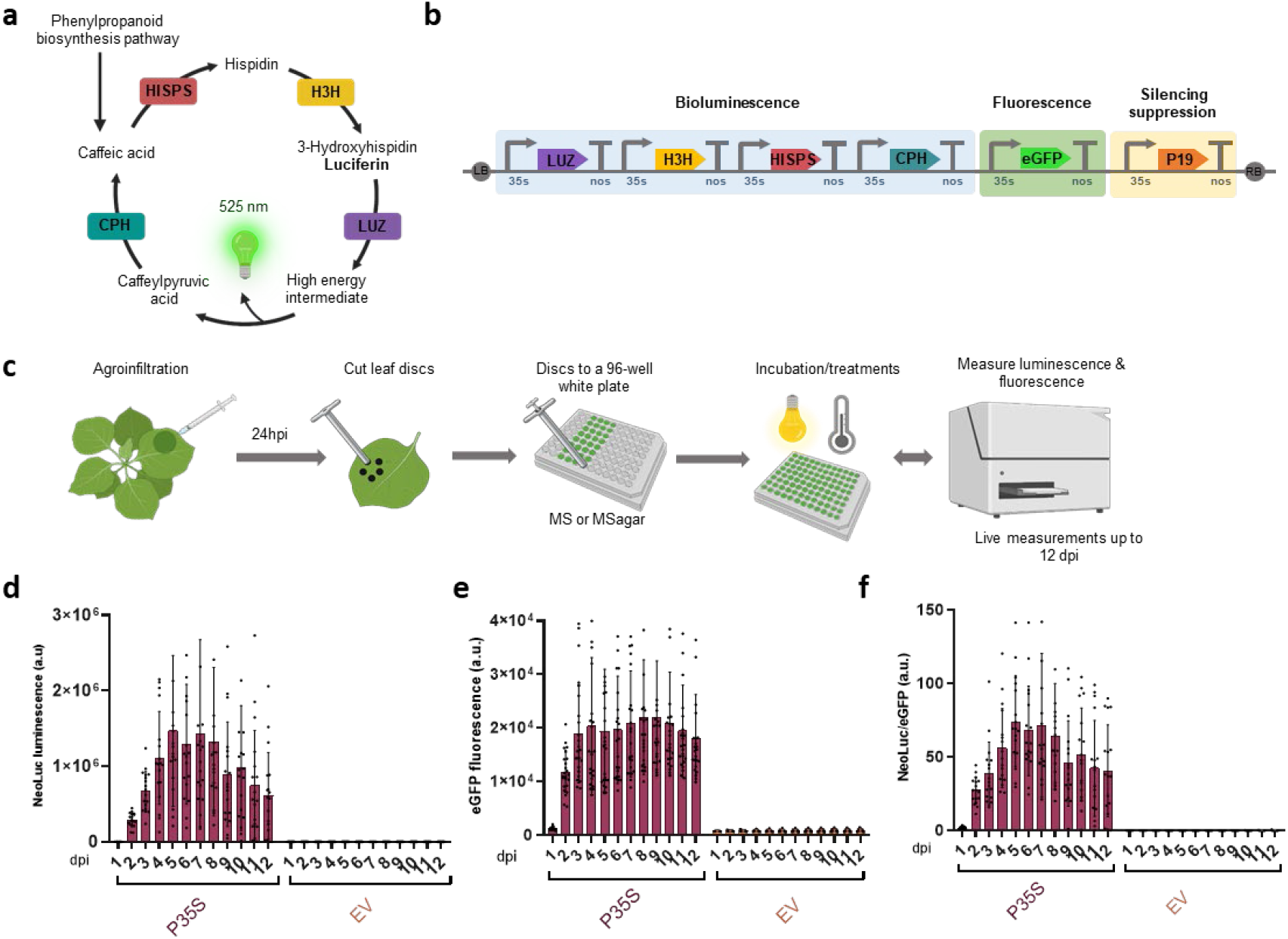
A quantitative bioluminescence reporter system for *N. benthamiana*. (**a**) Schematic representation of the *N. nambi* fungal bioluminescence pathway (FBP). (**b**) Scheme of T-DNA construct used for the dual reporter system, including the transcriptional units for the constitutive expression of the FBP genes (*Luz, H3H, HispS, CPH*), the enhanced GFP gene (*eGFP*) and the *p19* silencing suppressor gene. (**c**) Schematic representation of the experimental procedure used for luminescence and fluorescence measurements. (**d**) NeoLuc signal (a.u.) and (**e**) eGFP fluorescence signal (a.u.) of *N. benthamiana* leaf discs transiently expressing the reporter module depicted in (b) (P35S) or an empty vector (EV). (**f**) NeoLuc/eGFP ratios calculated with the luminescence and fluorescence values plotted in (d) and (e). Error bars indicate SD (n=24).

Here we report the development of a wide dynamic range, high-throughput reporter system for fast gene circuit testing based on FBP. This system combines fungal luminescence as model circuit actuator with eGFP as normalizing signal and employs transiently transformed *N. benthamiana* leaf disks as *in vivo* chassis. A key improvement in this new system is the employ of *HispS* gene, the enzyme catalyzing the first committed step in the pathway, as the gene that connects the reporter module with the upstream elements in the circuit. The use of *HispS* instead of *Luz* results in a wider dynamic range of the reporter, up to levels equivalent to the FLuc/RLuc system. The simultaneous measurements of LUZ-driven fungal luminescence (NeoLuc) and green fluorescence (eGFP) showed no interference using standard equipment. By employing this new methodology, we rapidly characterized several SGCs involving hormone and optogenetic sensors. Furthermore, we show the optimization of new copper switches and the *de novo* development of a dual-input control circuit for metabolic pathways.

## RESULTS

### Setting up an easy-to-use quantitative bioluminescence reporter system

To establish the basis for a dual reporter system, we first assembled a T-DNA containing the transcriptional units (TUs) for the 35SCaMV-driven constitutive expression of the four genes of the pathway (*Luz, H3H, CPH, HispS*) together with the silencing suppressor P19 and an enhanced GFP (eGFP) reporter (Fig. 1b). When agroinfiltrated in *N. benthamiana* leaves, eGFP showed a strong normalizing signal with minimal interference with *N. nambi* LUZ emission and therefore this construct was employed in the search for high throughput experimental procedures that minimized sample manipulation while producing consistent NeoLuc and eGFP measurements. We opted for a process consisting in the agroinfiltration of the FBP/eGFP module in intact *N. benthamiana* leaves followed, after 24h, by the excision and transfer of the leaf discs to 96-well plates containing sucrose-free MS medium (Fig. 1c). Next, dual NeoLuc/eGFP measurements were performed in plates from 1 dpi onwards using standard luminometer equipment. Leaf disks showed measurable expression levels up to 12 dpi (Fig 1d-f). Based on these results, we built a FBP/eGFP reporter module, initially setting the *Luz* gene under the control of the query promoter, as previously described by Khakhar et al., (2020). We compared then the performance of this initial configuration (referred as FBP[*Luz*]/eGFP), with the standard dual FLuc/RLuc system employing five constitutive promoters covering a wide range (∼150 fold) of relative transcriptional activities (RTAs). In the standard FLuc/RLuc system, RTA is calculated by normalizing FLuc/RLuc output ratios of a problem circuit with those provided by a constitutive Nopaline Synthase promoter (PNOS) in pre-defined standard experimental conditions (see Fig. 2a). To facilitate comparisons between the two reporter systems, we calculated for each promoter the area under the curve (AUC) of the NeoLuc/eGFP values measured through the time course of the experiment (Fig. 2b), and normalized this value against the AUC generated by the PNOS promoter, obtaining the final FBP[*Luz*]-RTA values. This parameter was thereafter employed to represent the overall transcriptional output of each SGCs. Unfortunately, as can be observed by comparing Fig. 2a and 2c, there was a clear saturation in the FBP[*Luz*]-RTA values that severely affected stronger promoters, thus limiting the applicability of the reporter system in its current configuration.

**Figure 2.**
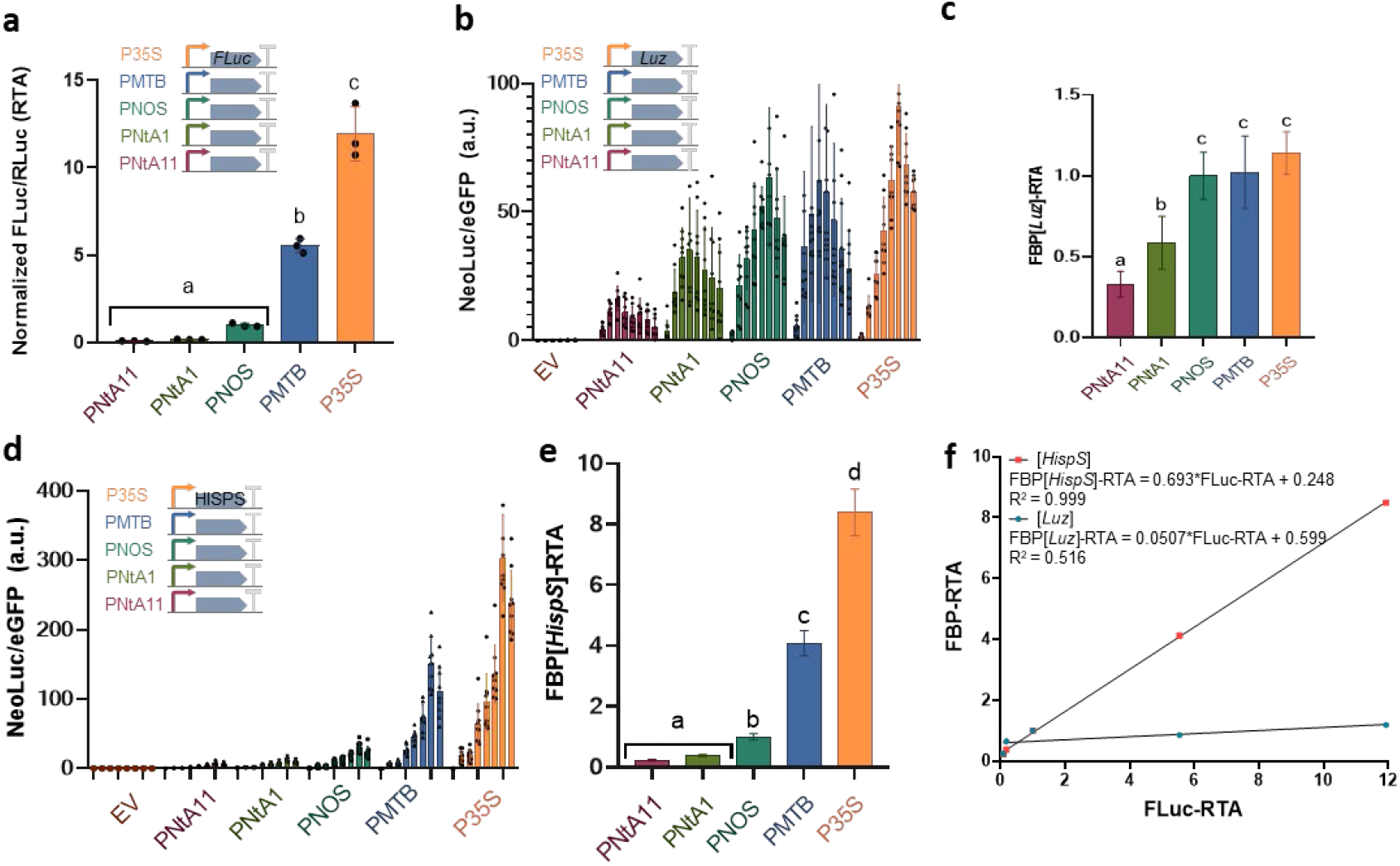
Optimization of the FBP/eGFP reporter employing five constitutive promoters. (**a**) Relative transcriptional activities (RTA) of four different constitutive promoters (PNtA11, PNtA1, PMTB and P35S) transiently expressed in *N. benthamiana*, calculated as FLuc/RLuc ratios for each promoter and normalized with the FLuc/RLuc ratios conferred by the constitutive PNOS promoter. (**b**) NeoLuc/eGFP ratios and (**c**) FBP[*Luz*]-RTA values of *N. benthamiana* leaf discs transiently expressing the FBP reporter with the *Luz* gene driven by four different constitutive promoters. (**d**) NeoLuc/eGFP ratios and (**e**) FBP[*HispS*]-RTA values of *N. benthamiana* leaf discs transiently expressing the FBP reporter with *HispS* gene driven by four constitutive promoters. (**f**) Correlation of normalized FLuc/RLuc ratios (FLuc-RTA) depicted in (a) with FBP[*Luz*]-RTA and FBP[*HispS*]-RTA values depicted in (c) and (e), respectively. An empty vector (EV) was used as a negative control in all experiments. Error bars indicate SD (a, b and d) and SE (c and e): n=9. Statistical analyses were performed using One-way ANOVA (Tukey’s multiple comparisons test, P-Value ≤ 0.05). Variables within the same statistical groups are marked with the same letters.

We reasoned that the observed signal saturation could be due to the choice of *Luz* as the connection point for the transcriptional output, as we earlier determined that the *HispS* gene and to a lesser extend the *H3H* gene, are the transcriptionally limiting steps in the pathway^9^ (see Supplementary Fig. 1). We therefore changed the reporter conformation, setting the query promoter position driving *HispS* expression instead of *Luz* (referred to as FBP[*HispS*]/eGFP configuration). Using this new setup, signal saturation was eliminated for stronger promoters (Fig. 2d), thus generating FBP[*HispS*]-RTA values that showed a strong lineal correlation (R^2^= 0.9997) to the classical RTA data observed with FLuc/RLuc system, also for strong promoters (Fig. 2e, 2f). In general, we observed that FBP[*HispS*]-RTA estimations are higher than FLuc/RLuc for weak promoters (PNTA11, PNTA1), probably reflecting a higher sensitivity, but somehow lower for stronger promoters, possibly due to some remaining signal saturation. Given the strong lineal correlation between the two systems, a conversion factor of 0.7 could be applied as a rule of thumb for comparisons of strong promoters if required. Although the sensitivity of FBP[*HispS*]-RTA in the *N. benthamiana* agroinfiltration system seems sufficient to cover the whole range of promoter strengths usually employed in PSB, we also assayed the introduction of a fifth enzyme, NPGA, known to activate HISPS^8^. The co-transformation with NPGA significantly increased the NeoLuc/eGFP values for all promoters assayed (Supplementary Fig. 2), offering additional sensitivity, if required.

### Phytohormone sensors with the FBP[HispS]/eGFP reporter

As a first immediate application of the new reporter, we designed new circuits to detect changes in phytohormone concentrations in the incubation media. A first device was set up to respond to increasing ABA concentrations using the plant endogenous hormone receptor components and an ABA-responsive promoter (p*MAPKKK18)* driving the expression of *HispS* (Fig. 3a). As shown (Fig. 3b, 3c), this simple sensor detected the presence of ABA with high confidence at concentrations above 25 μM. A second, more sophisticated and fully synthetic phytohormone detection system was next established for auxin sensing. In this design, we employed an anti-Cas protein, AcrIIA4, fused to the auxin degron mAID. The mAID:AcrIIA4 fusion functioned as a negative regulator of the CRISPR-based transcriptional activation (CRISPRa) mediated by the previously described dCasEV2.1 programmable transcription factor^10^. dCasEV2.1 strongly activates *HispS* by binding the PDFR promoter that contains a target site for a specific gRNA (gDFR) (Fig. 3d). dCasEV2.1 activity is strongly repressed by mAID:AcrIIA4, but in the presence of IAA, mAID:AcrIIA4 is degraded, releasing the activation of the transcriptional signal. As shown in Fig. 3e and Fig. 3f, a concentration of 0.1 μM IAA in the medium is sufficient to produce significant changes in FBP[*HispS*]-RTAs values. We also determined that the excised leaf disks retain the capacity to respond to the IAA up to 2 dpi (Fig. 3g and Supplementary Fig. 3). At later time points, the system’s responsiveness to the hormone is sharply lost.

**Figure 3.**
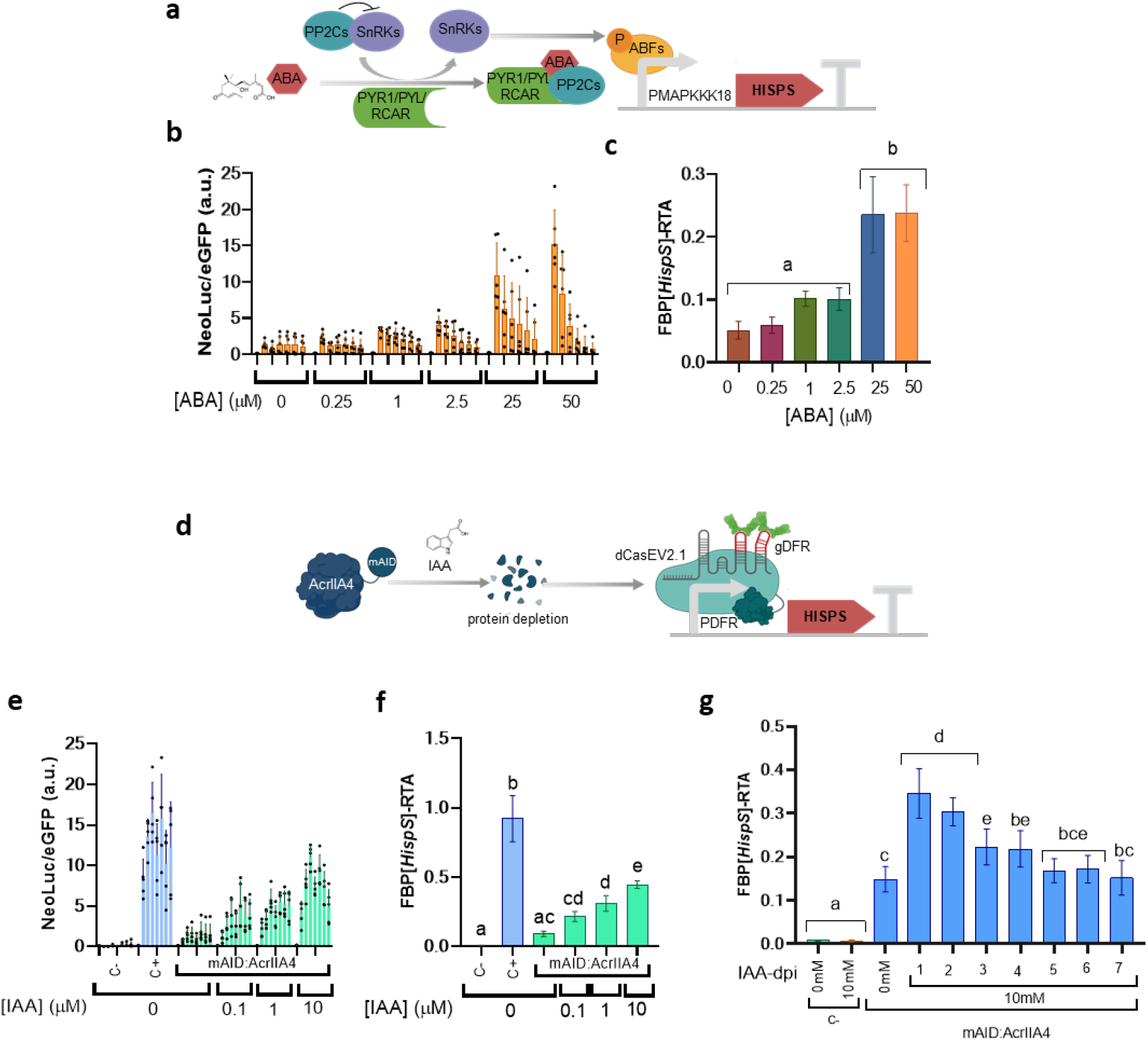
Phytohormone sensors with the FBP[*HispS*]/eGFP reporter. (**a**) Schematic representation of *HispS* gene activation driven by the ABA inducible promoter PMAPKKK18 (ABA sensor). (b) NeoLuc/eGFP ratios and (**c**) FBP[*HispS*]-RTA values of the ABA sensor in *N. benthamiana* leaf discs treated with different ABA concentrations. (**d**) Schematic representation of an IAA mediated AcrIIA4 protein depletion switch for the regulation of the dCasEV2.1-mediated reporter activation. (**e**) NeoLuc/eGFP ratios and (**f**) FBP[*HispS*]-RTA values of the IAA-mediated depletion switch in *N. benthamiana* leaves expressing dCasEV2.1 with an unspecific gRNA (C-) or the specific gRNA gDFR without (C+) or with mAID:AcrIIA4 treated with different IAA concentrations. (**g**) FBP[*HispS*]-RTA values of the IAA-mediated depletion switch in *N. benthamiana* leaves expressing dCasEV2.1 with an unspecific gRNA (C-) or the specific gRNA gDFR with mAID:AcrIIA4. Samples were either left untreated (0mM), or treated with 10 mM IAA at different time points. Error bars indicate SD (b and e) and SE (c, f and g): n=6 for (b, c and g) and n=5 for (e, f). Statistical analyses were performed using One-way ANOVA (Tukey’s multiple comparisons test, P-Value ≤ 0.05). Variables within the same statistical groups are marked with the same letters.

### Diversifying copper switches in plants using the FBP/eGFP reporter system

Repurposing agrochemicals such as copper sulfate into signaling molecules is an attractive option in PSB since agrochemicals can be realistically applied to genetically engineered crops at large scale. A *Saccharomyces cerevisiae* copper sensor was recently optimized in our group^11^. This copper sensor is based on the copper-responsive transcription factor CUP2. In the presence of Cu^2+^, CUP2 specifically binds the so-called Copper Binding Site (CBS) operator producing downstream gene activation mediated by a GAL4 transcriptional activation domain (GAl4AD) (Fig. 4a). We decided to further characterize different configurations of a copper switch taking advantage of the new data-rich FBP[*HispS*]/eGFP reporter. In its simplest configuration, the copper-responsive processor comprised a synthetic promoter containing CBS and the minimal promoter of the DFR gene (mDFR). As reflected in Figure 4b and Supplementary Fig. 4a, copper-dependent transcriptional activation was detected for this circuit at all assayed concentrations (from 0.1 to 5.0 mM), however the best activations were observed at the lowest concentration, suggesting certain toxicity effects at higher concentrations, something consistent with the lower levels of fluorescence recorded for these samples (Supplementary Fig. 4b). The system responsiveness to copper was maintained for 4dpi, although after the first 24h responsiveness declines sharply (Fig. 4c and Supplementary Fig. 4c). The second configuration assayed comprised an intermediary step involving dCasEV2.1 activation. The involvement of dCasEV2.1 (CRISPRa) provides two potential advantages to the copper switch. On one hand it enables multiplexing of downstream targets, and on the other hand, it offers additional control points, since the expression of all three components comprising the programmable activator (the gRNA, dCas9:EDLL and the MS2:VPR fusion, see Fig. 4d) can be regulated, jointly or independently, by one or more triggers. To reinforce copper control of the switch, we introduced a configuration where both the dCas9:EDLL and the MS2:VPR are regulated by copper (Fig. 4d). This regulation is important because it has been earlier observed that the introduction of intermediary CRISPRa elements raise background expression levels as side effect^11^. As can be observed in Fig. 4e, by introducing the new two copper control points, we obtained a CRISPRa-mediated copper switch with negligible background expression levels and with a dose-response curve similar to that exhibited by the simple (not multiplexable) switch (see also Supplementary Fig. 4d).

**Figure 4.**
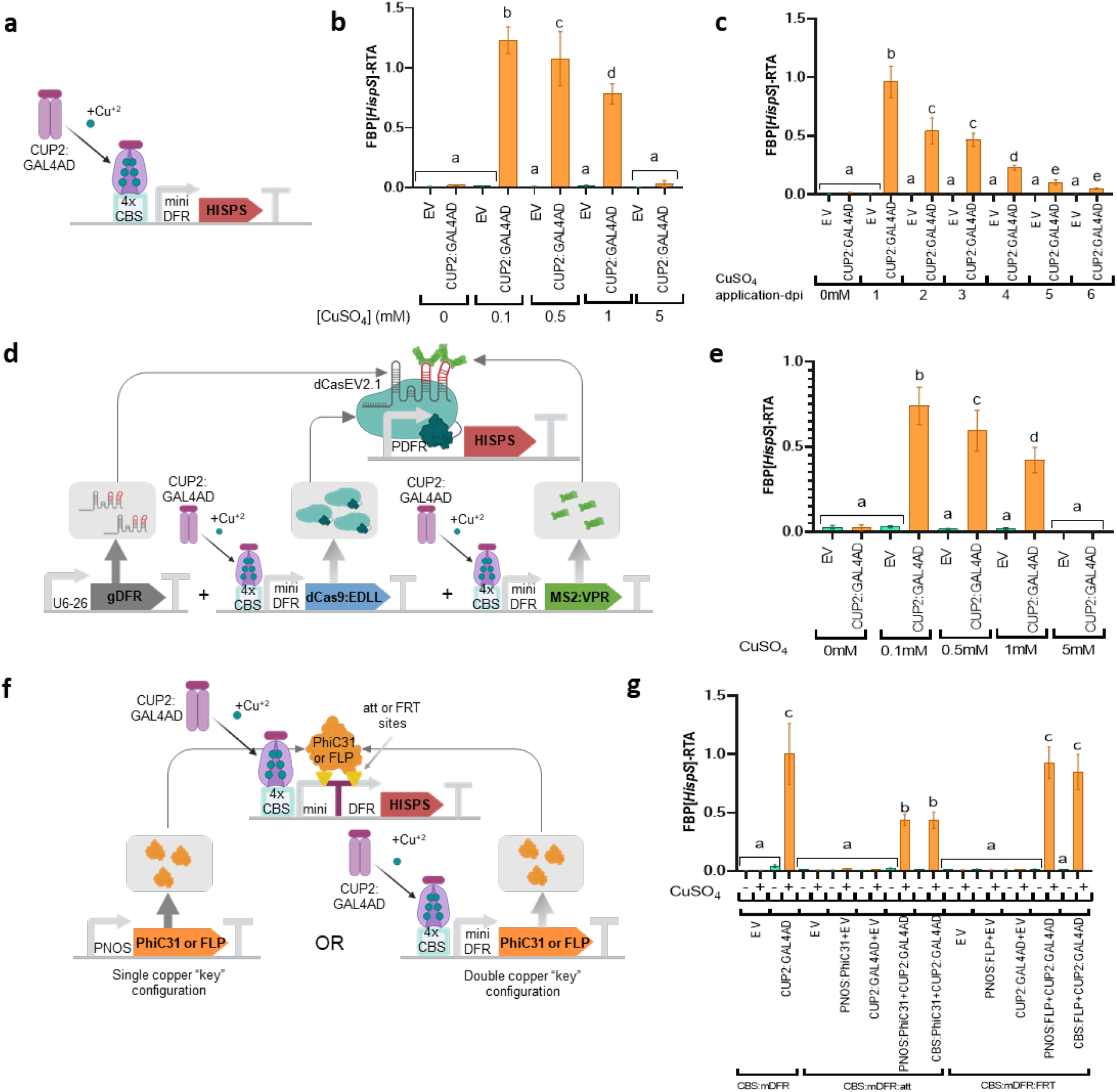
Alternative designs for a copper switch assayed with the FBP[*HispS*]/eGFP reporter. (**a**) Schematic representation of *HispS* gene activation driven by the copper inducible transcription factor CUP2:GAL4AD. (**b**) FBP[*HispS*]-RTA values of the simple copper switch depicted in (a) treated with different CuSO4 concentrations or (**c**) incubating with 0.1 mM CuSO4 at different time points. EV represents control experiments where the sensor CUP2:GAL4AD was substituted by an empty vector. (**d**) Schematic representation of *HispS* gene activation driven by a copper inducible dCasEV2.1. In this circuit both protein components of dCasEV2.1 (dCas9:EDLL and MS2:VPR) are regulated by copper, while the gRNA (gDFR) is constitutively expressed. (**e**) FBP[*HispS*]-RTA values of the copper swich represented in (d) with or without CUP2:GAL4AD and treated with different CuSO4 concentrations. (**f**) Schematic representation of *HispS* gene activation controlled by a recombinase (FLP or PhiC31) and a copper sensor. In this design the minimal DFR promoter is disrupted by the octopine synthase terminator flanked by recombination sites (FRT or att). The presence of both components (CUP2:GAL4AD and recombinase) is required for *HispS* expression. (**g**) FBP[*HispS*]-RTA values of the copper switch depicted in (f) with or without CUP2:GAL4AD and the corresponding recombinase (FLP or PhiC31) either driven by a PNOS or by the controlled by copper (CBS). Samples were treated with 0.1 mM CuSO4 (+) or let untreated (-). EV represents an empty vector. Error bars indicate SE: n=9 for (b) and (e); n=6 for (c) and (g). Statistical analyses were performed using One-way ANOVA (Tukey’s multiple comparisons test, P-Value ≤ 0.05). Variables within the same statistical groups are marked with the same letters.

Finally, we used FBP[*HispS*]/eGFP to study a new copper switch in which we introduced an “expression lock” mediated by a phage recombinase. For this, we inserted a terminator (OCS) within the mDFR promoter driving *HispS* expression, thus interrupting any residual activity conferred by the upstream CBS operator. The terminator was flanked by site-specific recombinase sites and therefore it was removable upon the presence of the recombinase, following a design earlier proposed by Lloyd et al.^12^. As depicted in Fig. 4f, we generated two alternative versions of the OCS-mDFR “locked” promoter, containing recombination sites for either the PhiC31 Integrase (att sites) or the *S. cerevisiae* Flipasse (FLP), (FRT sites). The copper sensing module comprising the CUP2:Gal4 transactivator was functionally linked to the FBP/eGFP actuator through the “locked” promoter and was co-agroinfiltrated either with a constitutively expressed recombinase (single copper “key” configuration) or with a copper-controlled one (double “key” configuration). As shown in Fig. 4g and Supplementary Fig. 4e, only when all elements of the circuit are present, and in the occurrence of copper sulfate, bioluminescence is produced. The FLP single and double “key” circuits produced output RTAs comparable to those produced by the simplest “unlocked” circuit, with lower background levels on average. Single and double key configurations of the PhiC31 version of the lock also maintained low background levels but the overall RTA levels in the presence of copper were significatively lower. For both types of SGCs, no significant differences were observed between single and double key configurations.

### Dual chemical-optogenetic control of the FBP

We next decided to explore new control mechanisms in plants that integrate two independent inputs, namely a chemical and an optogenetic signal. Rather than designing an AND logic gate as described elsewhere^13,14^, we took advantage of the fact that FBP is a metabolic pathway to assay an *in-series* control mechanism where each input signal controls a different step in the pathway. First, we built and assayed two different optogenetic control circuits. The blue light (BL) sensor, assayed in plants for the first time, relies on the interaction of *A. thaliana* CRY2 and CIB1 proteins with BL. By linking the *E. coli* Etr DNA-binding domain (EBD) to one of the proteins (CRY2:EBD) and an activation domain to the second interactor (CIB1:VPR), activation of FBP[*HispS*] only occurs in the presence of BL (Fig. 5a). Similarly, in the red-light (RL) sensor earlier characterized in plants^15^, PIF6:EBD and PHYB:VP16 interact upon illumination with RL, activating transcription of downstream genes (Fig. 5b). Both circuits were assayed in two alternative configurations, each one using a different minimal promoter upstream the *HispS* gene (mDFR or mCMV). As can be observed when comparing Fig. 5c, 5d with Fig. 5e, 5f, the behavior of the RL circuit was superior to the BL both in terms of background expression and in terms of maximum expression levels. In both systems, the employ of the minimal DFR promoter resulted in larger induction rates. The remarkable performance of the RL/mDFR circuit prompted us to employ it in combination with the single copper switch described in Fig. 4a to create the final double-input auto-bioluminescent circuit. To this end, we generated two FBP reporter modules containing (i) the *HispS* gene driven by the copper responsive promoter and (ii) the *H3H* gene driven by the RL responsive promoter, and vice versa (Fig. 5g). *N. benthamiana* plants were then co-infiltrated with agrobacterium cultures containing the FBP reporter module plus either one or both sensing modules (PIF-PHYB for RL or CUP2 for copper). Excised discs were subsequently placed onto MS 96-well plates with or without 0.1 mM CuSO_4_ and incubated in red light or in darkness. As observed in Fig. 5h and 5i, bioluminescence was only detected in the presence of both sensor modules and in RL/copper combined treatments. In all other circumstances, autonomous bioluminescence remained in an OFF state, with very low basal levels. Configuration II, linking *H3H* with copper and *HispS* with RL modules respectively, resulted in significantly higher RTA levels, while maintaining similar background expression in single-input treatments.

**Figure 5.**
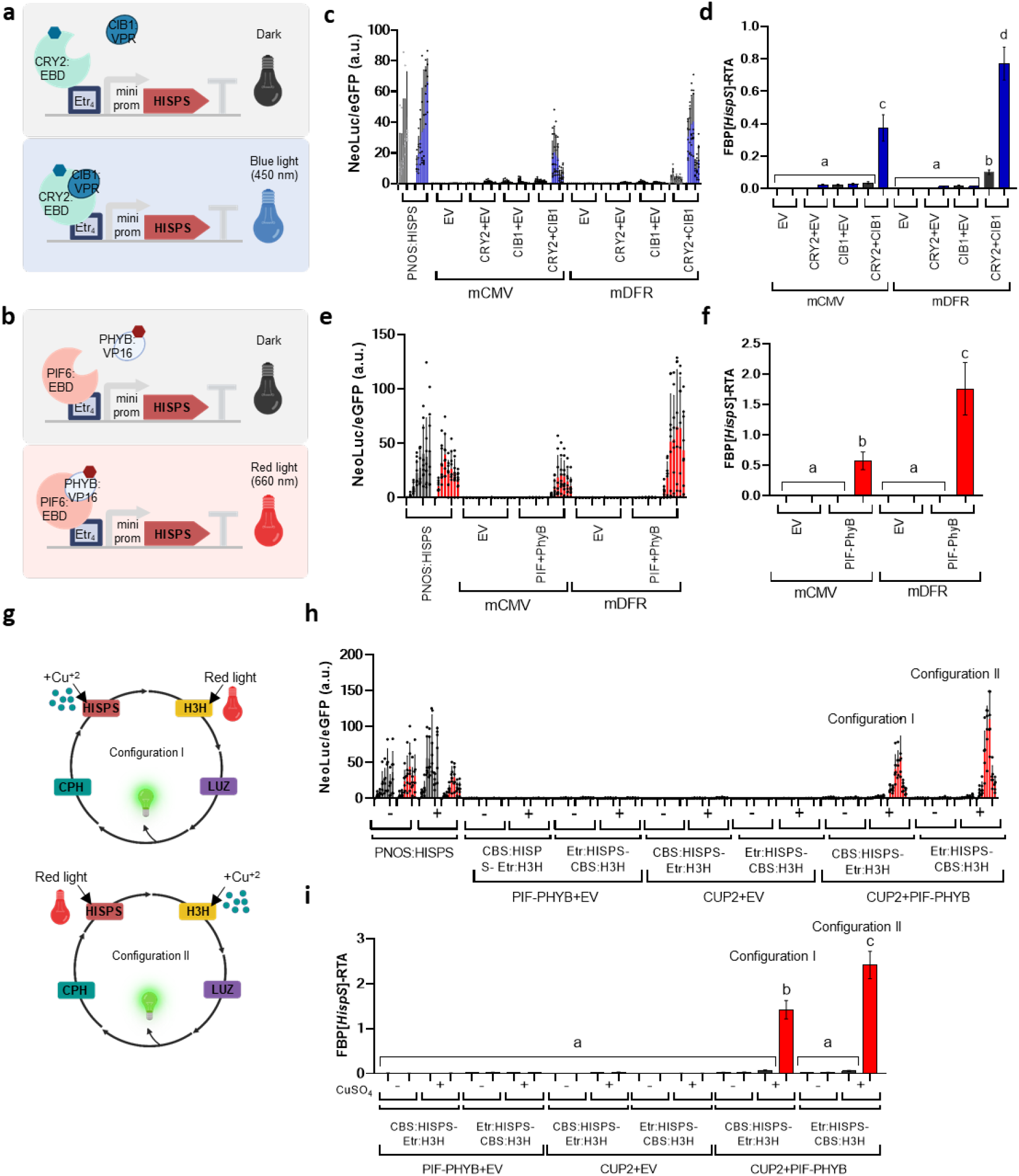
Dual optogenetic and chemical control of the fungal bioluminescence pathway (FBP). (**a**) Schematic representation of the CRY2:EBD-CIB1:VPR blue light (BL)-dependent *HispS* activation. (**b**) Schematic representation of the PIF6:EBD-PHYB:VP16 red light (RL)-dependent *HispS* activation. (**c**) NeoLuc/eGFP ratios and (**d**) FBP[*HispS*]-RTA values of the BL switch depicted in (a) expressed in *N. benthamiana* leaves, with or without CRY2:EBD and CIB1:VPR. Two minimal promoters driving *HispS* were assayed: mDFR and mCMV. Blue bars are BL treatments; black bars are dark treatments. (**e**) NeoLuc/eGFP ratios and (**f**) FBP[*HispS*]-RTA values of the RL switch depicted in (b) assayed in *N. benthamiana*, with or without PIF6:EBD and PHYB:VP16. Red bars are RL treatments; black bars are dark treatments. (**g**) Schematic representation of the two alternative configurations assayed for the dual RL and copper control of the FBP. (**h**) NeoLuc/eGFP ratios and (**i**) FBP[*HispS*]-RTA values of the circuits depicted in (g) in Configurations I and II. (-) are water treatments and (+) represents 0.1 mM CuSO4 treatments. CBS:HISPS-Etr:H3H represents Configuration I of the circuit. Etr:HISPS-CBS:H3H represents Configuration II. CUP2+PIF-PHYB are samples containing the entire circuit. PIF-PHYB+EV and CUP2+EV represent control samples were one of the sensor modules was missing and substituted by an empty vector. PNOS:HISPS is a normalizing control. EV samples were agroinfiltrated with an empty vector. Error bars indicate SD (c, e and h) and SE (d, f and i): n=9 for (c), (d), (e) and (f) and n=6 for (h) and (i)). Statistical analyses were performed using One-way ANOVA (Tukey’s multiple comparisons test, P-Value ≤ 0.05). Variables within the same statistical groups are marked with the same letters.

### Generation of stable reporter plant lines for FBP[HispS]/eGFP

In the pursuit of reporter system simplification, we decided to generate stable transgenic *N. benthamiana* plants carrying three (*H3H, Luz* and *CPH*, named as FBPΔHispS) (Fig 6a) or two (*CPH* and *Luz*, named as FBPΔHispSΔH3H) (Fig 6b) of the four genes of the *N. nambi* luminescence metabolic pathway. Transformed plants were screened by transiently expressing the missing pathway genes plus the eGFP. As observed in Figures 6c and 6d, several positive plants were recovered in which both NeoLuc and eGFP were readily detected. NeoLuc/eGFP levels were up to ten times higher in the FBPΔHispSΔH3H configuration, which could be related to the number of T-DNA copies per cell delivered via agroinfiltration. We next calibrated the FBPΔHispS line #1 (which delivered the best luminescence levels) by comparing NeoLuc/eGFP readings of P35S and PNOS promoters driving *HispS*. As observed in Figure 6e and 6f, the P35S/PNOS ratios in *HispS*-complemented FBPΔHispS plants were very similar to those observed with the same promoters when all FBP genes are delivered transiently (P35s RTA = 7.62+/-0.78 for all-transient system and 7.19+/-1.06 for FBPΔHispS plants). This indicates that the stable integration of the final enzymes in the pathway is a suitable strategy to alleviate the cargo of the transiently expressed modules while preserving the dynamic range of the reporter system.

**Figure 6.**
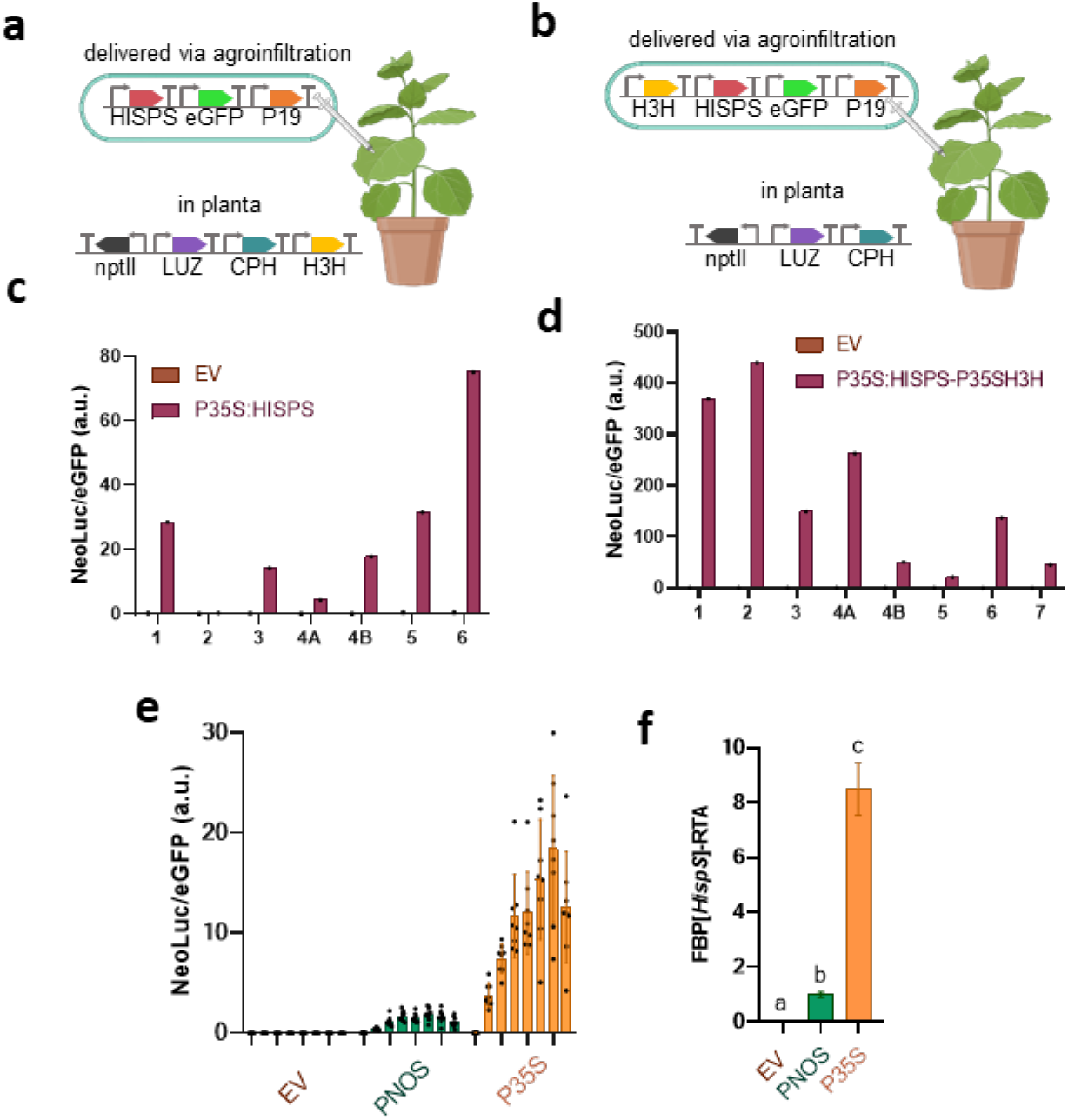
*N. benthamiana* plants expressing components of the FBP reduce the size of the plasmids delivered via agroinfiltration. Schematic representation of the FBP genes incorporated in the plant and those delivered via agroinfiltration in (**a**) the [−HISPS] plants and (**b**) the [−HISPS, −H3H] plants. NeoLuc/eGFP ratios of (**c**) seven independent T0 [−HISPS] and (**d**) eight independent T0 [−HISPS, −H3H] *N. benthamiana* plants transiently expressing either an empty vector (EV) or the missing components of the FBP reporter in each plant. (**e**) NeoLuc/eGFP ratios and (**f**) FBP[*HispS*]-RTA values of T0 [−HISPS] line 1 transiently expressing either an EV or the *HispS* gene driven by the PNOS or P35S promoters. Error bars indicate SD (e) and SE (f): n=9. Statistical analyses were performed using One-way ANOVA (Tukey’s multiple comparisons test, P-Value ≤ 0.05). Variables within the same statistical groups are marked with the same letters.

## DISCUSSION

One of the engineering principles in Synthetic Biology consists in advancing innovations through the Design–Build–Test–Learn (DBTL) cycle, a recursive loop used to develop new synthetic biological devices. Plant genetic engineers are progressively adopting DBTL paradigm despite the many plant-specific experimental obstacles. In the last decade, building DNA constructs was the main bottleneck in the cycle, however modular cloning methods have enormously facilitated multigene SGCs building, moving the constrains to the testing steps. Here, we have developed a new auto-bioluminescence/fluorescence reporter system that can significantly contribute to bypass this second DBTL bottleneck. We demonstrate that, by linking the circuit transcriptional outputs to *HispS*, and by including eGFP as normalizer, the FBP/eGFP reporter overcomes the limitations of previously described FBP reporters^8^ and serves as a reliable and highly sensitive alternative for FLuc/RLuc in SGC characterization.

A potential constrain for the use of FBP as reporter is the dependence on the plant’s endogenous metabolic status, in particular the intracellular concentration of caffeic acid. Depletion of the cell’s caffeic acid pool is unlikely, as FBP is a cyclic pathway and this metabolite is recycled. However, fluctuations in substrate availability, or uncharacterized feedback mechanisms may occur, and will need to be considered in experimental designs. Actually, substrate availability issues are not exclusive of FBP, as differential tissue permeability also affects luminescent reporters in other *in vivo* systems where the substrate luciferin needs to be added exogenously^16^. However as shown extensively in this work, the interferences of the plant’s metabolic status, if any, can be easily neutralized by a careful standardization of the testing procedures in the *N. benthamiana* agroinfiltration process, as proposed earlier^5^. *N. benthamiana* leaf agroinfiltration has gained great popularity among plant biotechnologists, and it is also the system of choice for plant-based bioproduction at industrial level^17^. Furthermore, the rich multi-omic resources made available recently completes the requirements for *N. benthamiana* to become a standard chassis for fast DBTL engineering^18,19^, without precluding the adaptation of FBP/eGFP to other experimental systems as tobacco/*Arabidopsis* protoplasts or seedlings^20^.

In the experiments described here we have generated a total of 1500 experimental points to characterize nine different plant SGCs, employing significantly less effort and much less resources than with any previous reporter system. For convenience, we have used AUC to condense the time-course data points in a single RTA value. We consider this parameter much more informative of promoter/circuit transcriptional activity than single time-point measurements usually offered in FLuc/RLuc experiments. Furthermore, by recording circuit behavior as time-course, any abnormal behavior attributed, for instance, to circadian clock interferences, can be easily accounted for and eventually corrected. Some of the SGCs described here recapitulated previous designs as proof of the performance of FBP/eGFP^10,11,21^, whereas others are new to plants. Among them, we have developed new circuit configurations for agrochemical (copper) control of gene expression with direct application in crops, and we have designed a new dual optogenetic/agrochemical control system that could be applied to the tight regulation of metabolic pathways in sensible metabolic engineering approaches. The expanded testing potential of the FBP/eGFP reporter is one example of the many new opportunities that the long-time pursued autonomous luminescence can offer to plant biotechnology.

## MATERIALS AND METHODS

### GoldenBraid cloning

DNA constructs used in this work were assembled using GoldenBraid (GB)^22^. Coding sequences of the *N. nambi* bioluminescence genes *Luz, H3H, CPH* and *HispS* previously codon optimized for expression in *N. tabacum* by Mitiouchkina et al.^7^ were ordered as gBlocks™ from IDT and domesticated at www.gbcloning.upv.es/do/domestication/. Level 0, Level 1 and Level>1 assemblies were performed using GB as previously described^22^. Level 0 assemblies were confirmed by restriction analysis and Sanger sequencing and Level 1 and Level >1 assemblies were verified by restriction analysis. All plasmids used in this work are listed in Supplementary Table 1.

### N. benthamiana transient expression

For transient expression assays, plasmids were transferred to *A. tumefaciens* strain GV3101 or EHA105 (see Supplementary Table 1 for details). Five to six weeks old *N. benthamiana* plants grown at 24°C and 16 h (light)/ 20°C and 8 h (darkness) conditions were used. For agroinfiltration, overnight *A. tumefaciens* cultures were pelleted and resuspended in agroinfiltration buffer (10 mM MES, pH 5.6, 10 mM MgCl2 and 200 μM acetosyringone) to an optical density of 0.1 at 600 nm (OD_600_). Bacterial suspensions were incubated for 2 h at room temperature on a horizontal rolling mixer, and then mixed for co-expression experiments, in which more than one GB element was used. Finally, agroinfiltrations were carried out through the abaxial surface of the three youngest fully expanded leaves of each plant with a 1-ml needle-free syringe. Detailed information of the experimental design can be found in the Supplementary Methods section.

### NeoLuc luminescence and eGFP fluorescence measurements and calculation of relative transcriptional activity

Leaf disc samples were collected at 24 hours post infiltration using a cork borer with a diameter of 0.5 cm. Two or three discs per agroinfiltrated leaf were excised. Leaf discs were transferred to a white 96-well microplate containing sucrose-free MS agar (or liquid) and incubated at 16 h light/8 h dark, 25°C, 60–70% humidity, 250 μmol m^−2^ s^−1^ photons. Luminescence and fluorescence were recorded on each leaf disc every 24 h (from 1 to 8-12 dpi). All measurements were carried out with a GloMax^®^-Multi Detection System (Promega). *N. nambi* luciferase (NeoLuc) activity was determined with the luminescence module using an integration time of 10s. eGFP fluorescence was determined using the blue optical kit (Ex: 490nm, Em: 510-570nm). NeoLuc/eGFP ratios were calculated for each leaf disc at each timepoint. The FBP[*Luz*]-RTA and FBP[*HispS*]-RTA values were calculated by dividing the area under the curve (AUC) of the NeoLuc/eGFP ratios produced by each sample from 2 to 8 dpi by the AUC produced by a PNOS reference calculated in an equivalent manner and assayed in parallel. The AUC were calculated using GraphPad with the default parameters. GB4148 (PNOS:Luz) was used for normalization of data displayed in Fig. 2c, GB4807 (PNOS:HISPS) was used for normalization of data displayed in Fig. 6g and GB4401 (PNOS:HISPS) was used for normalization of the rest of the experiments in this work.

### FLuc and RLuc activity assays and calculation of relative transcriptional activity

For the FLuc/RLuc assay, leaf samples were collected at 4 dpi. One 0.8 cm diameter disc per agroinfiltrated leaf was excised using a cork borer. Leaf discs were frozen in liquid nitrogen and subsequently homogenized. The Dual-Glo® Luciferase Assay System (Promega) was used for sample analysis. Briefly, 180 μl of Passive Lysis buffer were added to the homogenized sample, vortexed and centrifuged for 15 min at 14000 x*g* at 4°C. Crude extract (10 μl) was mixed with 40 μl of LARII and FLuc activity was determined using a GloMax^®^-Multi Detection System (Promega) with a 2 s delay and a 10 s integration times. After the measurements, 40 μl of Stop&Glo reagent were added per sample and RLuc activity was determined using the same protocol. Sample FLuc/RLuc ratios were calculated as the average value of the three independent agroinfiltrated leaves. Relative transcriptional activities (RTAs) were calculated as previously described^23^. Briefly, the FLuc/RLuc ratios in each sample normalized with the FLuc/RLuc ratios produced by a PNOS reference (GB1116) assayed in parallel.

### Sample size and statistical analysis

The sample number and statistical analysis considerations for each experiment are indicated in the corresponding figure. For FLuc/RLuc assays, 3 biological replicates at a single timepoint were analysed. For NeoLuc/eGFP determination, at least 5 biological replicates and 7 to 11 timepoints were analysed. Statistical analyses were performed using One-way ANOVA (Tukey’s multiple comparisons test, P-Value ≤ 0.05). Plotting and statistic tests were all performed with GraphPad.

### N. benthamiana stable transformation

For *N. benthamiana* stable transformation, *A. tumefaciens* LBA4404 was used. Transformation was performed following a standard protocol. Briefly, fully expanded leaves of *N. benthamiana* were sterilized with 5% commercial bleach (40 g of active chlorine per liter) for 10 min followed by four consecutive washing steps with sterile demi-water. Leaf discs (d = 0.8 cm) were cut with a cork borer and incubated overnight in co-culture plates [4.9 g/L Murashige and Skoog medium (MS) supplemented with vitamins (Duchefa), 3% sucrose (Sigma-Aldrich), 0.9% Phytoagar (Duchefa), 1 mg/L BAP (Sigma-Aldrich), 0.1 mg/L NAA (Sigma-Aldrich), pH 5.7]. Leaf discs were incubated for 15 min with the Agrobacterium culture (OD600 = 0.3). Then, the discs were returned to the co-culture plates and incubated for 2 days in darkness. Next, discs were transferred to selection medium [4.9 g/L MS supplemented with vitamins (Duchefa), 3% sucrose (Sigma-Aldrich), 0.9% Phytoagar (Duchefa), 1 mg/L BAP (Sigma-Aldrich), 0.1 mg/L NAA (Sigma-Aldrich), 500 mg/L carbenicillin, 100 mg/L kanamycin, pH 5.7]. Discs were transferred to fresh medium every seven days until shoots appeared (4–6 weeks). Shoots were cut and transferred to rooting medium [4.9 g/L MS supplemented with vitamins (Duchefa), 3% sucrose (Sigma-Aldrich), 0.9% Phytoagar (Duchefa), 500 mg/L carbenicillin, 100 mg/L kanamycin, pH 5.7] until roots appeared. Growing conditions were in all steps 16 h light/8 h dark, 25°C, 60–70% humidity, 250 μmol m−2 s−1 photons.

## SUPPORTING INFORMATION

**Supplementary Figure 1**. Analysis of the dynamic range of different combinatorial configurations of the FBP.

**Supplementary Figure 2**. NPGA enhances luminescence production allowing increased sensitivity of the FBP[*HispS*] reporter.

**Supplementary Figure 3**. Phytohormone sensors with the FBP[HispS]/eGFP reporter.

**Supplementary Figure 4**. Alternative designs for copper switches assayed with the FBP[HispS]/eGFP reporter.

**Supplementary Table 1**. List of GoldenBraid plasmids used in this work.

**Supplementary Methods**.

## ACKNOWLEDGEMENTS

This work was supported by grant PID2019-108203RB-I00 Plan Nacional I+D from the Ministerio de Ciencia e Innovación (Spain) through the Agencia Estatal de Investigación (co-financed by the European Regional Development Fund). C.C., M.V.-V. and E.M.-G are the recipients of fellowships FPI-UPV (PAID-01-20) from Universitat Politecnica de Valencia, APOSTD/2020/096 from the Generalitat Valenciana and FPU18/02019 from the Spanish Ministry of Science, Innovation and Universities, respectively. The authors would like to thank Matias Zurbriggen for sharing the plasmids of the red-light system and Jorge Lozano-Juste for sharing the plasmid with the *MAPKKK18* promoter. The authors also would like to thank Alberto Conejero for his assistance with data analysis.

## BIBLIOGRAPHY

1. Liu, W. & Stewart, C. N. Plant synthetic biology. Trends Plant Sci. 20, 309–317 (2015).

2. McCarthy, D. M. & Medford, J. I. Quantitative and Predictive Genetic Parts for Plant Synthetic Biology. Front. Plant Sci. 11, (2020).

3. Wang, Y.-H., Wei, K. Y. & Smolke, C. D. Synthetic biology: advancing the design of diverse genetic systems. Annu. Rev. Chem. Biomol. Eng. 4, 69–102 (2013).

4. Khosla, A., Rodriguez-Furlan, C., Kapoor, S., Van Norman, J. M. & Nelson, D. C. A series of dual-reporter vectors for ratiometric analysis of protein abundance in plants. Plant Direct 4, (2020).

5. Vazquez-Vilar, M. et al. GB3.0: a platform for plant bio-design that connects functional DNA elements with associated biological data. Nucleic Acids Res. 45, 2196– 2209 (2017).

6. Schaumberg, K. A. et al. Quantitative characterization of genetic parts and circuits for plant synthetic biology. Nat. Methods 13, 94–100 (2016).

7. Mitiouchkina, T. et al. Plants with genetically encoded autoluminescence. Nat. Biotechnol. 38, 944–946 (2020).

8. Khakhar, A. et al. Building customizable auto-luminescent luciferase-based reporters in plants. eLife 9, e52786 (2020).

9. Moreno-Giménez, E., Selma, S., Calvache, C. & Orzáez, D. GB_SynP: A Modular dCas9-Regulated Synthetic Promoter Collection for Fine-Tuned Recombinant Gene Expression in Plants. ACS Synth. Biol. 11, 3037–3048 (2022).

10. Selma, S. et al. Strong gene activation in plants with genome-wide specificity using a new orthogonal CRISPR/Cas9-based programmable transcriptional activator. Plant Biotechnol. J. 17, 1703–1705 (2019).

11. Garcia-Perez, E. et al. A copper switch for inducing CRISPR/Cas9-based transcriptional activation tightly regulates gene expression in Nicotiana benthamiana. BMC Biotechnol. 22, 12 (2022).

12. Lloyd, J. P. B. et al. Synthetic memory circuits for stable cell reprogramming in plants. Nat. Biotechnol. 40, 1862–1872 (2022).

13. Khan, M. A. et al. CRISPRi-based circuits for genetic computation in plants. http://biorxiv.org/lookup/doi/10.1101/2022.07.01.498372 (2022) xdoi:10.1101/022.07.01.498372.

14. Brophy, J. A. N. et al. Synthetic genetic circuits as a means of reprogramming plant roots. Science 377, 747–751 (2022).

15. Ochoa-Fernandez, R. et al. Optogenetic control of gene expression in plants in the presence of ambient white light. Nat. Methods 17, 717–725 (2020).

16. Shinde, R., Perkins, J. & Contag, C. H. Luciferin Derivatives for Enhanced in Vitro and in Vivo Bioluminescence Assays. Biochemistry 45, 11103–11112 (2006).

17. COVID vaccines grow on leaves. Nat. Biotechnol. 39, 649–649 (2021).

18. Ranawaka, B. et al. A multi-omic Nicotiana benthamiana resource for fundamental research and biotechnology. 022.12.30.521993 Preprint at https://doi.org/10.1101/2022.12.30.521993 (2022).

19. Kurotani, K.-I. et al. Genome Sequence and Analysis of Nicotiana benthamiana, the Model Plant for Interactions between Organisms. Plant Cell Physiol. pcac168 (2023) doi:10.1093/pcp/pcac168.

20. Furuhata, Y., Sakai, A., Murakami, T., Nagasaki, A. & Kato, Y. Bioluminescent imaging of Arabidopsis thaliana using an enhanced Nano-lantern luminescence reporter system. PLoS ONE 15, e0227477 (2020).

21. Calvache, C. et al. Strong and tunable anti-CRISPR/Cas activities in plants. Plant Biotechnol. J. (2021) doi:10.1111/pbi.13723.

22. Sarrion-Perdigones, A. et al. GoldenBraid 2.0: A Comprehensive DNA Assembly Framework for Plant Synthetic Biology. PLANT Physiol. 162, 1618–1631 (2013).

23. Vazquez-Vilar, M. et al. GB3.0: a platform for plant bio-design that connects functional DNA elements with associated biological data. Nucleic Acids Res. 45, 2196– 2209 (2017).

