## Supporting Information for "A quantitative autonomous bioluminescence reporter system with a wide dynamic range for Plant Synthetic Biology"

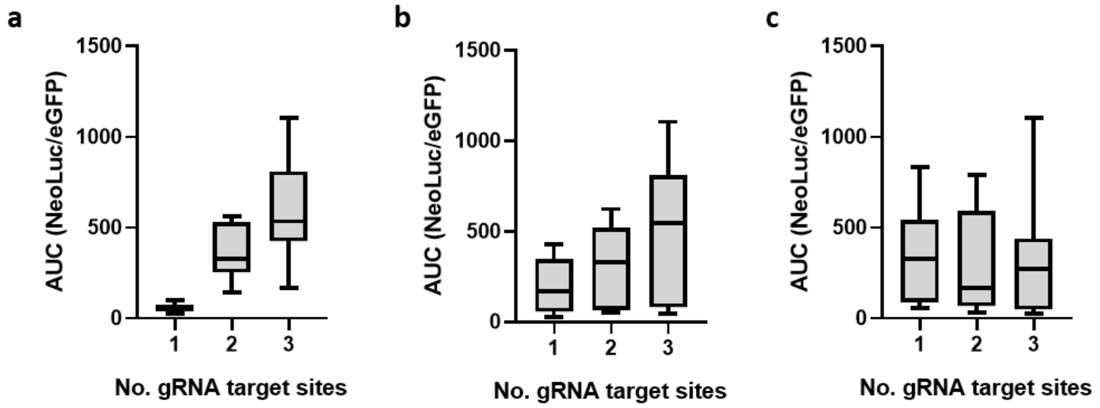


**Supplementary Figure 1. Analysis of the dynamic range of different combinatorial configurations of the FBP.** In each combination, CPH, eGFP and p19 are constitutively expressed with a 35S promoter, whereas Luz, H3H and HispS are driven by synthetic promoters containing one, two or three binding sites for the programable activator dCasEV2.1 summing up a total of ($3^3$ =27) configurations. Each plot represents the area under the curve (AUC) of NeoLuc/eGFP values for all 27 FBP reporter combinations when (**a**) Luz, (**b**) H3H and (**c**) HispS genes are expressed under synthetic promoters with either 1, 2 or 3 gRNA binding sites. For details see Moreno-Giménez et al.[1]

1. Moreno-Giménez, E., Selma, S., Calvache, C. & Orzáez, D. GB_SynP: a modular dCas9-regulated synthetic promoter collection for fine-tuned recombinant gene expression in plants. 2022.04.28.489949 Preprint at https://doi.org/10.1101/2022.04.28.489949 (2022).

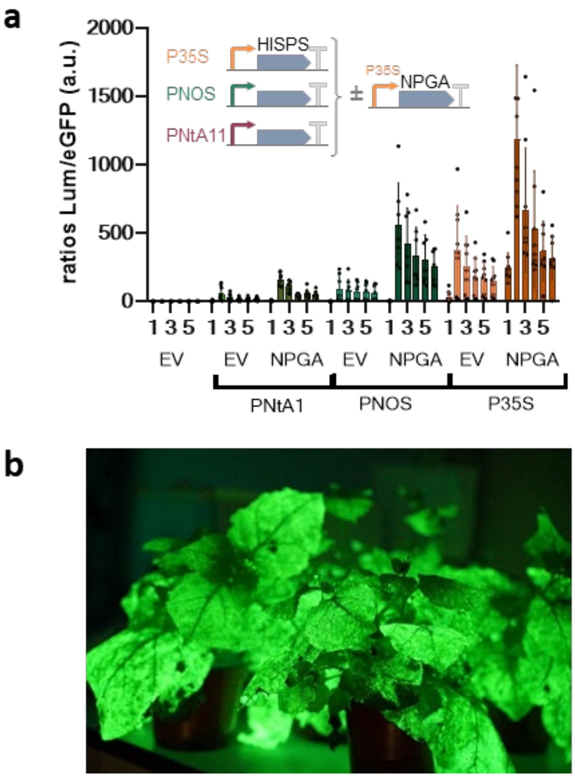


**Supplementary Figure 2. NPGA enhances luminescence production allowing increased sensitivity of the FBP[*HispS*] reporter.** (**a**) NeoLuc/eGFP ratios of *N.benthamiana* leaf discs transiently expressing the *HispS* gene driven by three different constitutive promoters and *H3H*, *Luz*, *CPH*, *eGFP* and *p19* with a 35S promoter, without (EV) or with NPGA. (**b**) *N. benthamiana* plants transiently expressing HISPS, H3H, LUZ, CPH, eGFP, P19 and NPGA. Images were taken with a reflex camera. An empty vector (EV) was used as a negative control. Error bars indicate SD (n=9).

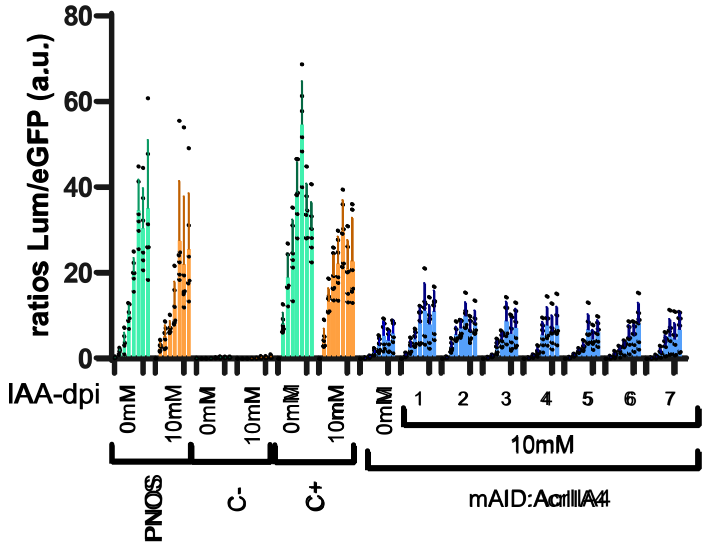


**Supplementary Figure 3. Phytohormone sensors with the FBP[*HispS*]/eGFP reporter.** NeoLuc/eGFP ratios of *N. benthamiana* leaves expressing the mDFR:*HispS* reporter, dCasEV2.1, the corresponding specific (C+) or unspecific (C-) gRNA and mAID:AcrIIA4 treated with 10mM IAA at different timepoints. Error bars indicate SD (n=5).

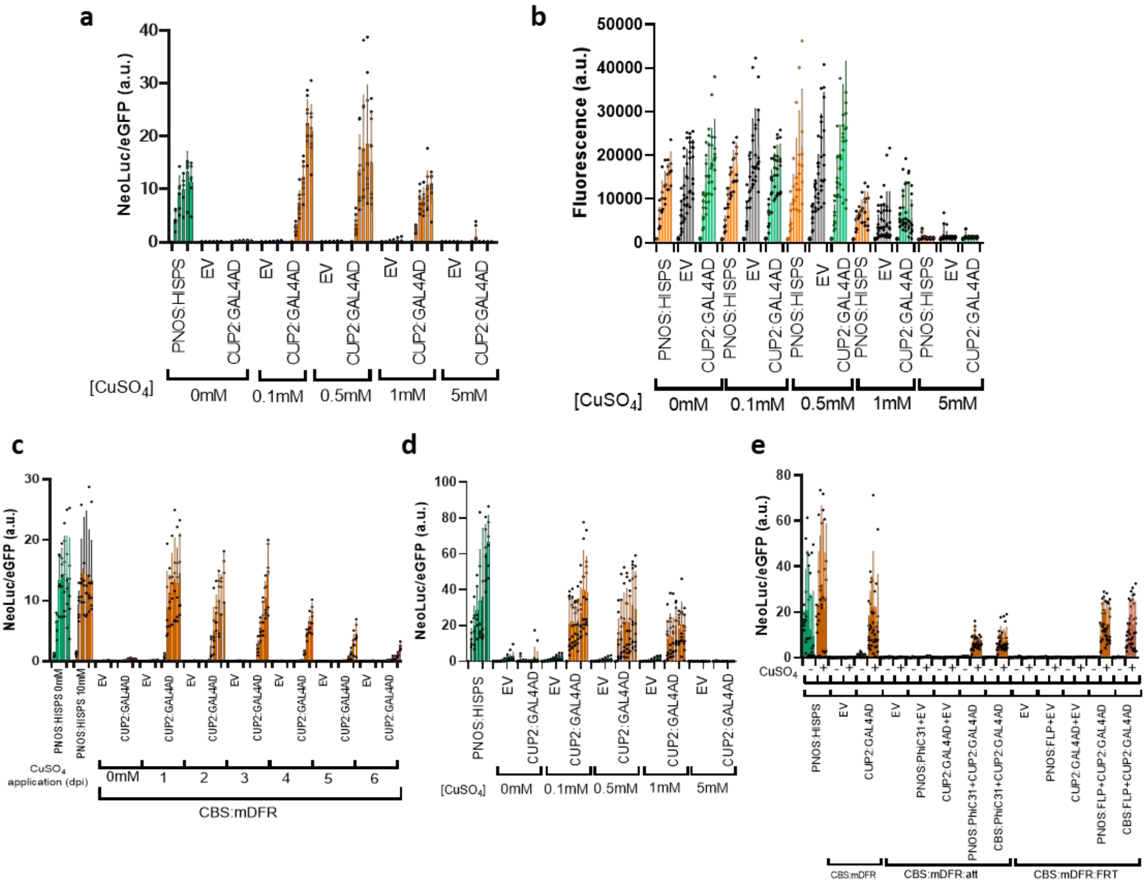


**Supplementary Figure 4. Alternative designs for copper switches assayed with the FBP[*HispS*]/eGFP reporter.** (**a**) NeoLuc/eGFP ratios and (**b**) fluorescence values of *N. benthamiana* leaf discs transiently expressing the *HispS* gene driven by the minimal DFR promoter preceded by a multiple copy of the CBS operator and constitutive *H3H*, *Luz*, *CPH*, *eGFP* and *p19*, with or without CUP2:GAL4AD and treated with different $CuSO_4$ concentrations. (**c**) NeoLuc/eGFP ratios of *N. benthamiana* leaf discs transiently expressing the *HispS* gene driven by the minimal DFR promoter preceded by a multiple copy of the CBS operator and constitutive *H3H*, *Luz*, *CPH*, *eGFP* and *p19*, with or without CUP2:GAL4AD treated with $CuSO_4$ at a final concentration of 0.1 mM at different timepoints. (**d**) NeoLuc/eGFP ratios of *N. benthamiana* leaf discs transiently expressing the *HispS* gene driven by the DFR promoter (including one gDFR binding site) and constitutive *H3H*, *Luz*, *CPH*, *eGFP* and *p19*, with or without CUP2:GAL4AD and treated with different $CuSO_4$ concentrations. (**e**) NeoLuc/eGFP ratios of *N. benthamiana* leaf discs transiently expressing the *HispS* gene driven by the minimal DFR promoter disrupted by the octopine synthase terminator flanked by recombination sites (FRT or att) and preceded by a multiple copy of the CBS operator and constitutive *H3H*, *Luz*, *CPH*, *eGFP* and *p19*, with or without CUP2:GAL4AD and the corresponding recombinase (FLP or PhiC31) either driven by a PNOS or by the CBS:mDFR promoter. $CuSO_4$ at a final concentration of 0.1 mM was applied. Error bars indicate SD: n=9 for (a), (b) and (d); n=6 for (c) and (e).

**Supplementary Table 1. List of GoldenBraid plasmids used in this work.** Sequence information is available at https://gbcloning.upv.es/search/ by entering the GB number.

| GB Number | Name | *Agrobacterium* strain used |
|---|---|---|
| GB1116 | pDGB3α1_PNOS:Luciferase:TNOS-SF-P35S:Renilla:TNOS-P35S:p19:TNOS | GV3101 |
| GB1119 | pDGB3α1_P35S:Luciferase:TNOS-SF-P35S:Renilla:TNOS-P35S:p19:TNOS | GV3101 |
| GB1236 | pDGB3α2_SF | GV3101 |
| GB1531 | pDGB3α1_PNos:PhiC31:TNos | GV3101 |
| GB1541 | pEGB_3alpha2 PNos:Cup2:Gal4AD:Tnos | GV3101 |
| GB1838 | pDGB3α1_U6-26-1gRNA-DFR F6x2 MS2scf | GV3101 |
| GB1203 | pDGB3α2_P35S:p19:TNOS | GV3101 |
| GB1399 | pDGB3α2_MTB:luc:Tnos-SF-35S:Ren:Tnos-35s:P19:Tnos-SF | GV3101 |
| GB2049 | pDGB3α1_U6-26-4gRNA Pnos-F6x2_35s-dCas9:EDLL-Tnos-U6-26-5gRNA Pnos scf F6x2 - 35s-Ms2:VPR-Tnos | GV3101 |
| GB2513 | pDGB3α1_dCas9EDLL-Ms2:VPR SF -gRNA DFR -150 2.1 | GV3101 |
| GB3668 | pDGB3α1_P35S:mAID:AcrIIA4:TNOS | GV3101 |
| GB3774 | pDGB3Ω1_CBS4:DFRmin:MS2:VPR:tNOS-CBS4:DFRmin:dCas9:EDLL:tNOS | GV3101 |
| GB4003 | pDGB3Ω1_PIF6-PhyB:VP16 | GV3101 |
| GB4080 | pDGB3Ω1_SF-EGFP-p19-BIoluc | EHA105 |
| GB4114 | pDGB3α1_PSlDFR:LUZ:TNOS-SF-H3H-HISPS-CPH-EGFP-p19 | EHA105 |
| GB4115 | pDGB3α1_Etr8:CMVmin:LUZ:TNOS-SF-H3H-HISPS-CPH-EGFP-p19 | EHA105 |
| GB4148 | pDGB3Ω1_PNOS:LUZ:TNOS-SF-SF-H3H-HISPS-CPH-EGFP-p19 | EHA105 |
| GB4149 | pDGB3Ω1_PMTB:LUZ:TNOS-SF-SF-H3H-HISPS-CPH-EGFP-p19 | EHA105 |
| GB4176 | pDGB3Ω1_PNtA1:LUZ:TNOS-SF-SF-H3H-HISPS-CPH-EGFP-p19 | EHA105 |
| GB4177 | pDGB3Ω1_PNtA11:LUZ:TNOS-SF-SF-H3H-HISPS-CPH-EGFP-p19 | EHA105 |
| GB4192 | pDGB3α1_P35S:EtrBD:CRY2PHR:TNOS | GV3101 |
| GB4193 | pDGB3α2_P35S:CIB1:VPR:TNOS | GV3101 |
| GB4216 | pDGB3Ω1_PNtA1:Luciferase:TNOS-SF-P35S:Renilla:TNOS-P35S:p19:TNOS | EHA105 |
| GB4217 | pDGB3Ω1_PNtA11:Luciferase:TNOS-SF-P35S:Renilla:TNOS-P35S:p19:TNOS | EHA105 |
| GB4401 | pDGB3Ω1_SF-PNOS:HISPS:TNOS-CPH-LUZ-H3H-EGFP-p19 | EHA105 |
| GB4402 | pDGB3Ω1_SF-PMTB:HISPS:TNOS-CPH-LUZ-H3H-EGFP-p19 | EHA105 |
| GB4403 | pDGB3Ω1_SF-PNtA1:HISPS:TNOS-CPH-LUZ-H3H-EGFP-p19 | EHA105 |
| GB4404 | pDGB3Ω1_SF-PNtA11:HISPS:TNOS-CPH-LUZ-H3H-EGFP-p19 | EHA105 |
| GB4540 | pDGB3Ω1_PDFR:HISPS-SF-CPH-SF-LUZ-H3H-EGFP-p19 | EHA105 |

| | | |
|---|---|---|
| GB4541 | pDGB3Ω1_Etr4:mDFR:HISPS-SF-CPH-SF-LUZ-H3H-EGFP-p19 | EHA105 |
| GB4542 | pDGB3Ω1_CBS4x:mDFR:HISPS-SF-CPH-SF-LUZ-H3H-EGFP-p19 | EHA105 |
| GB4544 | pDGB3Ω1_Etr8:CMVmin:HISPS-SF-CPH-SF-LUZ-H3H-EGFP-p19 | EHA105 |
| GB4405 | pDGB3Ω1_SF-MAR10-SF-nptII-CPH-LUZ-H3H-MAR10-SF | LBA4404 |
| GB4635 | pDGB3α1_CBS4x:mDFR:HISPS-Etr4:mDFR:H3HSF-LUZ-CPH-EGFP-p19 | EHA105 |
| GB4636 | pDGB3α1_Etr4:mDFR:HISPS-CBS4x:mDFR:H3HSF-LUZ-CPH-EGFP-p19 | EHA105 |
| GB4650 | pDGB3α2_ CBS:miniDFR:PhiC31:tNos | EHA105 |
| GB4652 | pDGB3α2_ CBS:miniDFR:FLP:TNOS | EHA105 |
| GB4653 | pDGB3α2_ pNos:FLP:tNos | EHA105 |
| GB4655 | pDGB3Ω1_CBS-att-OCS-HispS-tNos-SF-CPH-SF-LUZ-H3H-EGFP-p19 | EHA105 |
| GB4656 | pDGB3α1_ CBS-FRT-OCS-HispS-tNos-SF-CPH-SF-LUZ-H3H-EGFP-p19 | EHA105 |
| GB4665 | pDGB3Ω1_ MAR10-SF-nptII-LUZ-CPH-MAR10-SF | LBA4404 |
| GB4668 | pDGB3Ω1_pMAPKKK18:HISPS.TNOS-SF-CPH-SF-LUZ-H3H-EGFP-p19 | EHA105 |
| GB4807 | pDGB3Ω1_PNOS:HISPS-SF-EGFP-p19 | EHA105 |
| GB4808 | pDGB3Ω1_P35S:HISPS-SF-EGFP-p19 | EHA105 |
| GB4829 | pDGB3α1_P35S:NPGA:TNOS | EHA105 |
| GB4832 | pDGB3α1_HISPS-H3H-EGFP-p23 | EHA105 |

### Supplementary Methods

The tables provided below show the Agrobacterium cultures co-infiltrated in the same mix for each experiment.

*Assay for setting up a quantitative bioluminescence reporter system (Figure 1d, 1e, 1f)*

| *Construct* | EV | P35S |
|---|---|---|
| GB1236 | + | - |
| GB4080 | - | + |

*FLuc/RLuc constitutive promoter assay (Figure 2a)*

| *Construct* | EV | PNtA11 | PNtA1 | PNOS | PMTB | P35S |
|---|---|---|---|---|---|---|
| GB1236 | + | - | - | - | - | - |
| GB4217 | - | + | - | - | - | - |
| GB4216 | - | - | + | - | - | - |
| GB1116 | - | - | - | + | - | - |
| GB1399 | - | - | - | - | + | - |
| GB1119 | - | - | - | - | - | + |

*FBP[Luz]-RTA assay for constitutive promoters (Figure 2b, 2c)*

| *Construct* | EV | PNtA1 | PNtA1 | PNOS | PMTB | P35S |
|---|---|---|---|---|---|---|
| GB1236 | + | - | - | - | - | - |
| GB4177 | - | + | - | - | - | - |
| GB4176 | - | - | + | - | - | - |
| GB4148 | - | - | - | + | - | - |
| GB4149 | - | - | - | - | + | - |
| GB4080 | - | - | - | - | - | + |

*FBP[HispS]-RTA assay for constitutive promoters (Figure 2d, 2e)*

| *Construct* | EV | PNtA11 | PNtA1 | PNOS | PMTB | P35S |
|---|---|---|---|---|---|---|
| GB1236 | + | - | - | - | - | - |
| GB4404 | - | + | - | - | - | - |
| GB4403 | - | - | + | - | - | - |
| GB4401 | - | - | - | + | - | - |
| GB4402 | - | - | - | - | + | - |
| GB4080 | - | - | - | - | - | + |

*ABA-mediated regulation of HispS (Figure 3b, 3c)*

| *Construct* | 0µM | 0.25µM | 1µM | 2.5µM | 25µM | 50µM |
|---|---|---|---|---|---|---|
| GB4668 | + | + | + | + | + | + |

*Auxin Inducible Degron assay (Figure 3e, 3f)*

| ***Construct*** | **C- (0μM)** | **C+ (0μM)** | **mAID:AcrIIA4** | | | |
|---|---|---|---|---|---|---|
| | | | **0μM** | **0.25μM** | **1μM** | **2.5μM** |
| GB3668 | - | - | + | + | + | + |
| GB4540 | + | + | + | + | + | + |
| GB2513 | - | + | + | + | + | + |
| GB2049 | + | - | - | - | - | - |

*Auxin Inducible Degron time-course assay (Figure 3g and Supplementary Figure 3)*

| ***Construct*** | **0μM** | | | **10μM** | | | | | | | | | |
|---|---|---|---|---|---|---|---|---|---|---|---|---|---|
| | **PNOS** | **C-** | **C+** | **PNOS** | **C-** | **C+** | **mAID:AcrIIA4** | | | | | | |
| | | | | | | | **1dpi** | **2dpi** | **3dpi** | **4dpi** | **5dpi** | **6dpi** | **7dpi** |
| GB3668 | - | - | - | - | - | - | + | + | + | + | + | + | + |
| GB4540 | - | + | + | - | + | + | + | + | + | + | + | + | + |
| GB2513 | - | - | + | - | - | + | + | + | + | + | + | + | + |
| GB2049 | - | + | - | - | + | - | - | - | - | - | - | - | - |
| GB4401 | + | - | - | + | - | - | - | - | - | - | - | - | - |

*Copper-mediated regulation dose-response assay (Figure 4b and Supplementary Figure 4a, 4b)*

| ***Construct*** | **PNOS** | | | | | **EV** | | | | | **CUP2:GAL4AD** | | | | |
|---|---|---|---|---|---|---|---|---|---|---|---|---|---|---|---|
| | **0mM** | **0.1mM** | **0.5mM** | **1mM** | **5mM** | **0mM** | **0.1mM** | **0.5mM** | **1mM** | **5mM** | **0mM** | **0.1mM** | **0.5mM** | **1mM** | **5mM** |
| GB1236 | - | - | - | - | - | + | + | + | + | + | - | - | - | - | - |
| GB4542 | - | - | - | - | - | + | + | + | + | + | + | + | + | + | + |
| GB1541 | - | - | - | - | - | - | - | - | - | - | + | + | + | + | + |
| GB4401 | + | + | + | + | + | - | - | - | - | - | - | - | - | - | - |

*Copper-mediated regulation time-course assay (Figure 4c and Supplementary Figure 4c)*

| ***Construct*** | **PNOS** | | **CUP2:GAL4AD** | | **EV** | | | | | | **CUP2:GAL4AD** | | | | | |
|---|---|---|---|---|---|---|---|---|---|---|---|---|---|---|---|---|
| | **0mM** | **0.1mM** | **0mM** | **0.1mM** | **1dpi** | **2dpi** | **3dpi** | **4dpi** | **5dpi** | **6dpi** | **1dpi** | **2dpi** | **3dpi** | **4dpi** | **5dpi** | **6dpi** |
| GB1236 | - | - | - | - | + | + | + | + | + | + | - | - | - | - | - | - |
| GB4542 | - | - | + | + | + | + | + | + | + | + | + | + | + | + | + | + |
| GB1541 | - | - | + | + | - | - | - | - | - | - | + | + | + | + | + | + |
| GB4401 | + | + | - | - | - | - | - | - | - | - | - | - | - | - | - | - |

*Copper-activated dCas-mediated regulation assay (Figure 4e and Supplementary Figure 4d)*

| ***Construct*** | **PNOS** | **EV** | | | | | **CUP2:GAL4AD** | | | | |
|---|---|---|---|---|---|---|---|---|---|---|---|
| | **0mM** | **0mM** | **0.1mM** | **0.5mM** | **1mM** | **5mM** | **0mM** | **0.1mM** | **0.5mM** | **1mM** | **5mM** |
| GB1236 | - | + | + | + | + | + | - | - | - | - | - |
| GB4540 | - | + | + | + | + | + | + | + | + | + | + |
| GB1541 | - | - | - | - | - | - | + | + | + | + | + |
| GB3774 | - | + | + | + | + | + | + | + | + | + | + |
| GB1838 | - | + | + | + | + | + | + | + | + | + | + |
| GB4401 | + | - | - | - | - | - | - | - | - | - | - |

*Recombinase and copper control of FBP[HispS] (Figure 4g and Supplementary Figure 4e)*

| Construct | PNOS:HISPS | CBS:mDFR | | CBS:mDFR:att | | | | CBS:mDFR:FRT | | | |
|---|---|---|---|---|---|---|---|---|---|---|---|
| | | | | EV | | CUP2:GAL4AD | | EV | | CUP2:GAL4AD | |
| | | EV | CUP2:GAL4AD | PNOS:PhiC31 | CUP2:GAL4AD | PNOS:PhiC31 | CBS:PhiC31 | PNOS:FLP | CUP2:GAL4AD | PNOS:FLP | CBS:FLP |
| GB1236 | - | + | - | + | - | - | - | + | - | - | - |
| GB1541 | - | - | + | - | + | + | + | - | + | + | + |
| GB1531 | - | - | - | - | - | + | - | - | - | - | - |
| GB4650 | - | - | - | - | - | - | + | - | - | - | - |
| GB4652 | - | - | - | - | - | - | - | - | - | - | + |
| GB4653 | - | - | - | - | - | - | - | - | - | + | - |
| GB4542 | - | + | + | - | - | - | - | - | - | - | - |
| GB4655 | - | - | - | + | + | + | + | - | - | - | - |
| GB4656 | - | - | - | - | - | - | - | + | + | + | + |
| GB4401 | + | - | - | - | - | - | - | - | - | - | - |

*Blue-light system assay (Figure 5b, 5c)*

| Construct | PNOS:HISPS | CMVmin | | | | mDFR | | | |
|---|---|---|---|---|---|---|---|---|---|
| | | EV | CRY2+EV | CIB1+EV | CRY2+CIB1 | EV | CRY2+EV | CIB1+EV | CRY2+CIB1 |
| GB1236 | - | + | + | + | + | + | + | + | - |
| GB4401 | + | - | - | - | - | - | - | + | + |
| GB4192 | - | - | + | - | + | - | + | - | + |
| GB4193 | - | - | - | + | + | - | - | + | + |
| GB4544 | - | + | + | + | + | - | - | - | - |
| GB4541 | - | - | - | - | - | + | + | + | + |

*Red-light system assay (Figure 5e, 5f)*

| Construct | PNOS:HISPS | CMVmin | | mDFR | |
|---|---|---|---|---|---|
| | | EV | PIF+Phy | EV | PIF+Phy |
| GB1236 | - | + | + | + | - |
| GB4401 | + | - | - | - | - |
| GB4403 | - | - | + | - | + |
| GB4544 | - | + | + | - | - |
| GB4541 | - | - | - | + | + |

*Red-light and copper regulation of HispS and H3H (Figure 5h, 5i)*

| Construct | PNOS:HISPS | PIF-PhyB+EV | | CUP2:GAL4AD+EV | | PIF-PhyB+CUP2:GAL4AD | |
|---|---|---|---|---|---|---|---|
| | | CBS:HISPS-EtrH3H | Etr:HISPS-CBS:H3H | CBS:HISPS-EtrH3H | Etr:HISPS-CBS:H3H | CBS:HISPS-EtrH3H | Etr:HISPS-CBS:H3H |
| GB1236 | - | + | + | + | + | + | + |
| GB1541 | - | - | - | + | + | + | + |
| GB4401 | + | - | - | - | - | - | - |
| GB4403 | - | + | + | - | - | + | + |
| GB4635 | - | + | - | + | - | + | - |
| GB4636 | - | - | + | - | + | - | + |

*[ΔHISPS] plants screening assay (Figure 6c)*

| ***Construct*** | **EV** | **P35S: HISPS** |
|---|---|---|
| GB1236 | + | - |
| GB4808 | - | + |

*[ΔHISPS, ΔH3H] plants screening assay (Figure 6d)*

| ***Construct*** | **EV** | **P35S:HISPS-P35S:H3H** |
|---|---|---|
| GB1236 | + | - |
| GB4832 | - | + |

*[ΔHISPS] constitutive promoter assay (Figure 6e, 6f)*

| ***Construct*** | **EV** | **PNOS** | **P35S** |
|---|---|---|---|
| GB1236 | + | - | - |
| GB4807 | - | + | - |
| GB4808 | - | - | + |

*NPGA assay (Supplementary Figure 2)*

| ***Construct*** | **EV** | **PNtA1** | | **PNOS** | | **P35S** | |
|---|---|---|---|---|---|---|---|
| | | **EV** | **NPGA** | **EV** | **NPGA** | **EV** | **NPGA** |
| GB1236 | + | + | - | + | - | + | - |
| GB4403 | - | + | + | - | - | - | - |
| GB4401 | - | - | - | + | + | - | - |
| GB4080 | - | - | - | - | - | + | + |
| GB4832 | - | - | + | - | + | - | + |
